# Identification of Large Japanese field mouse *Apodemus speciosus* food plant resources in an industrial green space using DNA metabarcoding

**DOI:** 10.1101/2024.04.01.587655

**Authors:** Taichi Fujii, Hirokazu Kawamoto, Tomoyasu Shirako, Masatoshi Nakamura, Motoyasu Minami

**Author notes:** Corresponding author (TF). These authors contributed equally to this work.

## Abstract

DNA metabarcoding was employed to identify the food plant resources of the Large Japanese field mouse Apodemus speciosus, inhabiting an artificial green space on reclaimed land on the Chita Peninsula in Aichi Prefecture, Central Japan, from 2012 to 2014. DNA metabarcoding was performed using high-throughput sequencing of partial rbcL sequences extracted from feces samples collected in the study area. The obtained sequences, which were analyzed using a constructed local database, revealed that a total of 72 plant taxa were utilized as food plant resources by A. speciosus. Of these plant taxa, 43 could be assigned to species (59.7%), 16 to genus (22.2%), and 13 to family (18.1%). Of the 72 plant taxa identified in this study, the dominant families throughout all collection periods were Lauraceae (81.0% of 100 fecal samples), followed by Fagaceae (70.0%), Rosaceae (68.0%), and Oleaceae (48.0%). Fifty of the 72 plant taxa identified as food plant resources were woody plants. An analysis employing rarefaction techniques for each season in the study site indicated comprehensive coverage of the food plant resources, ranging from 86.4% in winter to 93.6% in spring. Further, 96.5% of the food plant taxa were found throughout the study period. The findings showed that DNA metabarcoding using a local database constructed from the National Center for Biotechnology Information (NCBI) database and field surveys was effective for identifying the dominant food plants in the diet of A. speciosus. The results of this study provided basic information that can be applied to formulation and implementation of management and conservation strategies for local wildlife.

## Introduction

Recently, artificial green spaces in urban areas have come to function as habitats for local wildlife and are considered to contribute positively towards regional biodiversity conservation. For example, species such as the red fox (*Vulpes vulpes*) [1–3], raccoon dog (*Nyctereutes procyonoides*) [2, 4], northern goshawk (*Accipiter gentilis*) [5], and Sunda scops-owl (*Otus lempiji*) [6–7] have been confirmed to inhabit and breed in artificial green spaces in urban areas. Considering that the wildlife in Japan has been threatened by extensive habitat destruction, mainly due to urbanization, artificial green spaces with habitats for small mammals in urban areas have contributed positively towards maintaining viable populations of wildlife [2, 8]. Residents of urban areas are aware of the value of artificial green spaces in urban areas for wildlife and wish to protect and coexist with wild animals [9]. Consequently, artificial green spaces in some areas are being assessed for their utility as wildlife habitats in Japan [2, 10]. In recent years, there has been a growing awareness of biodiversity conservation through the adoption of Other Effective Area-Based Conservation Measures (OECM) [11, 12], and the significance of managing green spaces effectively is increasing.

Of the small mammals in Japan, the Large Japanese field mouse *Apodemus speciosus* is an endemic and wild rodent that is widely distributed throughout the Japanese Archipelago, except the Ryukyu Islands at the southwestern-most tip of Japan [13]. While this species originally inhabited forest environments, it has also colonized artificially created green spaces in urban areas [2, 8]. In order to maintain the ecosystem health of the artificial green spaces in urban areas, it is necessary to maintain the populations of small mammals such as *A. speciosus*, which are preyed upon by carnivorous animals [14, 15]. Since *A. speciosus* populations are markedly affected by the abundance of food resources such as acorns [16], the preferential conservation of plant species that are utilized as food resources by this species is considered to be beneficial for the conservation of this rodent.

While several studies have examined the food plant resources of *A. speciosus* in the natural habitat [17–19], relatively few studies have examined food plant preference in artificial green spaces in urban environments [2]. Several methods have been employed to identify the food resources utilized by wildlife, including identification of the gastric or feces contents by dissection [20–23] and direct observations of foraging behavior [24, 25]. Since *A. speciosus* is a small rodent with a head and body length of 80–140 mm [13], identifying food resources based on direct observations of foraging behavior or by using fragmented plant residues in feces or in gastric contents is difficult [20, 26]. Recently, DNA metabarcoding using high-throughput sequencing (HTS) has been used to identify the diets of wildlife with higher sensitivity [18, 19, 27–38]. Several studies on the use of DNA metabarcoding to identify the food plant resources that are utilized by *A. speciosus* have been reported [18, 19]. However, the results obtained using the chloroplast *trn*L P6 loop intron region have yielded low accuracy in species-level estimates of food plant resources, with only 2.9% [18] and 15.7% [19] of detected plant taxa being accurately identified.

DNA metabarcoding has been shown to improve estimation accuracy by referencing local databases [38, 39]. Short sequence target regions, such as the chloroplast *trn*L P6 loop intron region used in a previous study [18], are often similar among genetically related plant species. Therefore, in order to improve the accuracy of identification at the species level, analyses based on the *rbcL* region that is longer than the *trn*L P6 loop intron region [40] will likely facilitate more accurate identification of the food resources utilized by *A. speciosus*. In this study, we identified *A. speciosus* food plant resources by DNA metabarcoding using a combination of the NCBI and a local database. The use of DNA metabarcoding is expected to provide a better understanding of the feeding habits of *A. speciosus* in urban environments. We consider that the results of this study will provide basic information that can be applied to the development of proper management and conservation strategies for local wildlife.

## Materials and methods

### Trapping rodents and fecal sample collection

The study site consists of industrial green spaces on reclaimed land on the Chita Peninsula in Aichi Prefecture, Central Japan (Fig 1). Since the industrial area opened in the 1970s, it has been separated from the surrounding area by Ise Bay to the west and Chita Industrial Road, which is more than 30 m wide, to the east. The vegetation at the site consists of a mixture of evergreen broad-leaved trees and deciduous broad-leaved trees, which were either planted when the industrial area was built or which naturally colonized the area from the surrounding environment. The dominant species in the study site were *Cinnamomum camphora*, *Lithocarpus edulis*, *Pterocarya stenoptera*, *Ligustrum lucidum*, and *Elaeocarpus zollingeri* var. *zollingeri*. The climate of the study area has a mean annual temperature of 16.1°C and precipitation of 1450.3 mm [41]. This study was conducted in full compliance with a license for capturing *A. speciosus* that was obtained from the Aichi Prefectural Government (ochi No. 3-1) and the Guidelines for Animal Treatment proposed by the Mammal Society of Japan, and all efforts were made to minimize suffering. We collected a total of 100 *A. speciosus* individuals (spring: 22 from Mar. to May, summer: 35 from Jun. to Aug, autumn: 32 from Sept. to Nov., winter: 11 from Dec. to Feb.) from the study site by Sherman-type live traps (6.5 × 5.5 × 16.0 cm, H. B. Sherman Traps, Tallahassee, FL, USA) from 2012 to 2014. The bait used for capturing *A. speciosus* consisted of a mixture of barley (*Hordeum vulgare* L. subsp. *vulgare*) and walnuts (*Juglans regia* L.) (15:1 w/w). Trapped individuals were released immediately after capture at the site of capture and all of the feces in the Sherman trap were collected for total DNA extraction. The fecal samples thus collected were stored at −20°C until total genomic DNA extraction.

**Fig 1.**
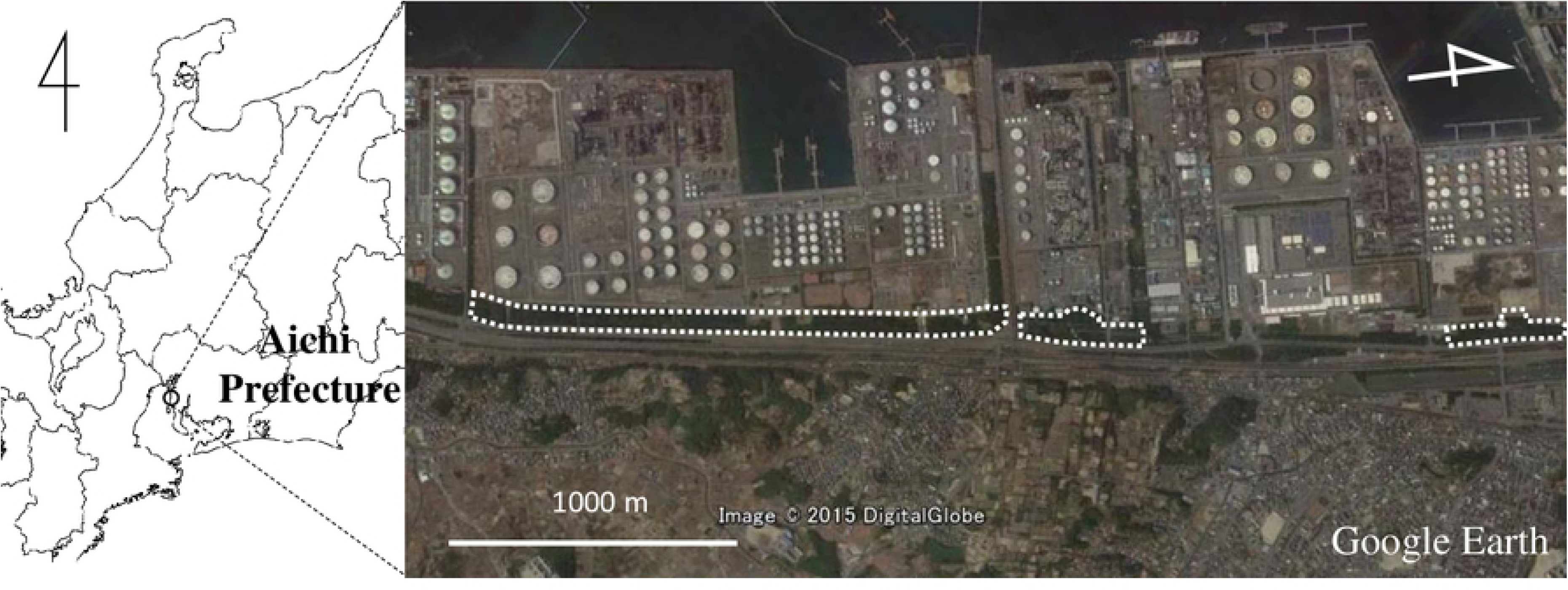
Map and aerial photograph of the industrial green spaces investigated in this study. Dotted lines indicate areas where the Large Japanese field mouse (*Apodemus speciosus*) was trapped. Map data were extracted from Google, DigitalGlobe.

### Partial sequencing of *rbcL* gene in fecal samples

Before performing DNA extraction, all fecal samples were dried overnight at 60°C. Total DNA was isolated from the dried feces (one fecal sample was ca. 60 mg) using a DNeasy Plant Mini Kit (Qiagen, Hilden, Germany) and purified using a Geneclean Spin Kit (MP-Biomedicals, Santa Ana, CA, USA). In this study, the identification of food plant species in feces was performed by DNA metabarcoding using HTS of the partial sequences of the *rbcL* region in chloroplast DNA (*rbcL* F3R3) [42]. Since amplicon sequencing by HTS generally decreases nucleotide diversity, adjacent DNA sequences on the flow cell can be misrecognized and low sequencing quality scores can result [43]. Therefore, in order to compensate for the reduction in nucleotide diversity, we used frame-shifting primers for the initial PCR. The initial PCR of *rbcL* F3R3 was performed in a reaction mixture of 12.0 μl with 6.0 μl of KAPA HiFi (Kapa Biosystems, Wilmington, MA, USA), 0.7 μl of 10 μM primer mix for *rbcL* F3R3 (S1 Table), 2.0 μl of template DNA, and 3.3 μl of dH_2_O under the following conditions: initial denaturation at 95°C for 3 min, followed by 35 cycles of denaturation at 98°C for 20 s, annealing at 56°C for 15 s, and extension at 72°C for 30 s, with a final extension step at 72°C for 5 min. The initial PCR products of *rbcL* F3R3 were purified using an Agencourt AMPure XP kit (Beckman Coulter, Fullerton, CA, USA). To construct the DNA libraries for the second PCR, we ligated the i7 and i5 indexes, which are required to identify each fecal sample, along with the P5 and P7 adapters (Illumina Inc.) necessary for Miseq sequencing, to the purified products of the initial PCR. The second PCR of *rbcL* F3R3 was performed in reaction mixtures of 24.0 μl containing 12.0 μl of KAPA HiFi, 2.8 μl of 10 μM forward and reverse index primers (S2 Table), 2.0 μl of 1/10 initial PCR products, and 4.4 μl of dH_2_O under the following conditions: initial denaturation at 95°C for 3 min, followed by 12 cycles of denaturation at 98°C for 20 s and extension at 72°C for 15 s, with a final extension step at 72°C for 5 min. The DNA libraries of *rbcL* F3R3 were purified using an Agencourt AMPure XP kit (Beckman Coulter) and sequenced using an MiSeq Reagent Kit v3 (600-cycle format; Illumina Inc.) on an Illumina MiSeq sequencer (Illumina Inc.) at IDEA Consultants, Inc. (Yaizu, Japan), following the manufacturer’s protocol.

### Sequence data analysis

Using the bcl2fastq v2.18 program (Illumina Inc.), the raw MiSeq data were first converted to FASTQ files, which were then demultiplexed using the clsplitseq function implemented in Claident [44]. In this study, the demultiplexed FASTQ files were analyzed using the amplicon sequence variant (ASV) method implemented in the DADA2 v1.10.1 package [45] using the R statistical software package [46]. As part of the quality filtering process, both forward and reverse sequences of *rbcL* F3R3 were trimmed to 200 bases. Trimming was performed based on visual inspection of the quality score distribution using the filterAndTrim function of DADA2 [45]. After trimming, forward and reverse sequences were combined using the mergePairs function of DADA2 [45], and chimeric DNA sequences were removed using the removeBimeraDenovo function of DADA2 [45]. Low-frequency ASVs, i.e., less than 1.0% of the total number of sequences, in each fecal sample were excluded from the DNA metabarcoding analysis. To normalize the number of DNA sequences in *rbcL* F3R3, rarefaction curves were calculated with the rarecurve function of vegan package ver. 2.5-5 [47] in R [46]. Using the calculated rarefaction curves, 1,000 *rbcL* F3R3 DNA sequences were normalized from each fecal sample using the rrarefy function of vegan R package [47]to determine the ASVs for plant species identification.

### Construction of partial *rbcL* database for the fecal samples collected in the study area

Based on vegetation surveys conducted by an environmental consulting company in 2001, and by us in this study, a total of 796 species were found in the study area (S3 Table). The *rbcL* F3R3 database of these 796 species, as well as barley (*H. vulgare* subsp. *vulgare*) and walnut (*J. regia*), which were used as bait for capturing mice, was constructed as follows. After obtaining the NCBI taxonomy IDs of the 798 species from the NCBI Taxonomy Database, we output the NCBI GI numbers containing the *rbcL* sequences of 798 species necessary for constructing a local database from the NCBI data using the taxonomy IDs of the 798 species and *rbcL* as keywords using the clretrievegi function implemented in Claident [44] (S3 Table). The NCBI GI numbers were then converted to GenBank format using the pgretrieveseq function implemented in Claident [44]. The fasta format data were converted from these GenBank format data using extractfeat the EMBOSS package [48]. Next, the sequences obtained were trimmed to match the *rbcL* F3R3 region (262 bp). As a result, within the 798 plant species, 147 were not registered in the NCBI (S3 Table). Furthermore, it is important to note that species identification for newly registered DNA sequences in the DNA databank is self-reported by the submitter. This can sometimes lead to sequences being associated with misidentified species [44–49]. To ensure the accurate identification of food plant resources for *A. speciosus* at this study site, we sequenced the *rbcL* F3R3 region of 104 predominant plant species at the study site using the following methods. Total DNA was extracted from plant specimens (ca. 0.8 cm^2^) using a DNeasy Plant Mini Kit (Qiagen) and purified using a Geneclean Spin Kit (MP-Biomedicals). PCR was then performed using a reaction mixture of 50 μl containing 1 unit of MightyAmp DNA Polymerase Ver. 2 (Takara, Shiga, Japan) and 0.32 µM of each primer according to the manufacturer’s instructions. The *rbcL* F3R3 amplicon was sequenced using the forward and reverse primers used in the aforementioned initial PCR for DNA metabarcoding using HTS to identify plant species in feces [42]. PCR amplification was performed using a DNA Thermal Cycler (GeneAmp PCR System 9700, Applied Biosystems, Foster City, CA) using an initial denaturation step of 98°C for 2 min, followed by 30 cycles of denaturation at 98°C for 10 s, annealing at 60°C for 15 s, and extension at 68°C for 20 s. All PCR products were purified using a QIAquick PCR Purification Kit (Qiagen) and subjected to dye-terminator cycle sequencing using DTCS Quick Start Mix (Beckman Coulter) and an automatic sequencer (CEQ 2000XL, Beckman Coulter).

Finally, the *rbcL* F3R3 sequences obtained from the NCBI database and the 104 plant species collected in the study site were combined to construct a local *rbcL* F3R3 database consisting of 651 plant species using the makeblastdb function implemented in NCBI’s BLAST+ software package [50] (S3 Table).

### Homology search and identification of food plant resources

DNA metabarcoding using HTS to identify the plant species in feces samples was conducted using a combination of our local *rbcL* F3R3 database and the NCBI database. Figure 2 shows a flowchart of the steps used to identify *A. speciosus* food plant resources in this study. To identify food plant species, homology searches were performed by comparing the ASVs obtained from each fecal sample against the local *rbcL* F3R3 database using the blastn function implemented in NCBI’s BLAST+ version 2.6.0+ package [50]. If the homology score from the local *rbcL* F3R3 database was less than 98%, these ASVs were subjected to a homology search using all of the published sequences deposited in the NCBI database. To ensure that the accuracy of the identification was sufficiently robust, DNA sequences for *rbcL* F3R3 with a homology of less than 98% in either database were excluded from further analysis. In addition, the ASVs that were identified as belonging to either barley (*H. vulgare* subsp. *vulgare*) or walnut (*J. regia*), which were used as bait, were also excluded from further analysis. In the event that a given sequence was identified as belonging to two or more taxa with the same score, that sequence was assigned to the highest taxonomic level that included both of those taxa. The accuracy of species identification was assessed by dividing the number of plant taxa identified to the species level by the total number of plant taxa. The ASV read abundance has been shown to be affected by plant-derived PCR inhibitors [51]. Therefore, ASV data were converted to presence/absence data.

**Fig 2.**
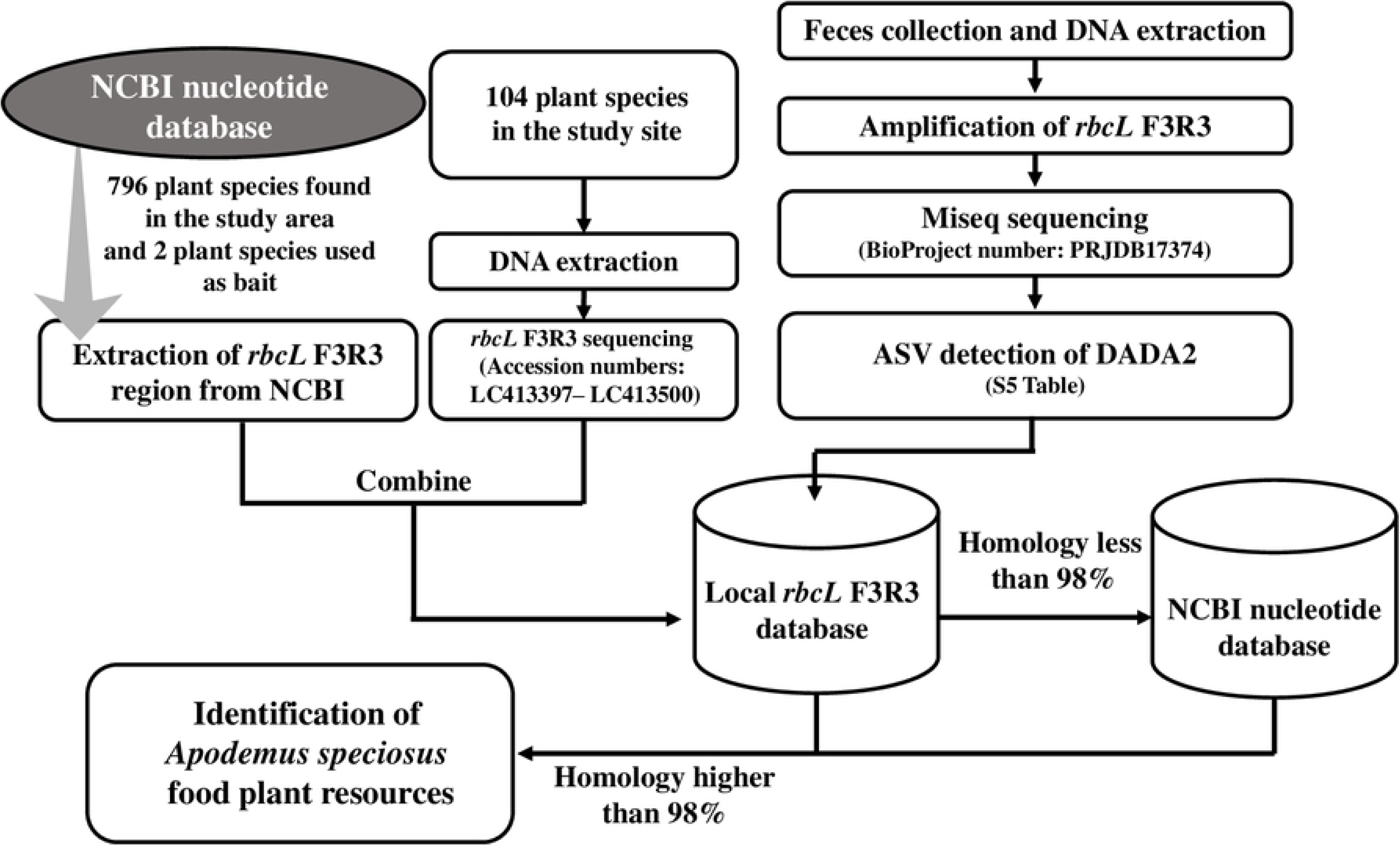
Flowchart showing the process used to identify food plant resources of Apodemus speciosus in this study.

## Results

### Construction of local *rbcL* F3R3 database

The 5934 DNA sequences in the local *rbcL* F3R3 database were obtained from 651 plant species in the NCBI database and 104 predominant plant species collected at the study site (S3 Table). The *rbcL* F3R3 sequences obtained from the 104 predominant plant species at the study site were registered with the DNA Data Bank of Japan under accession numbers LC413397–LC413500. In the local *rbcL* F3R3 database which contained 782 unique sequences, 631/782 (80.7%) were species-specific, 96/782 (12.3%) were the same at the genus level but not at the species level, 44/782 (5.6%) were the same at the family level but not at the genus level, and 11/782 (1.4%) could not be assigned even at the family level.

In this local *rbcL* F3R3 database, walnut (*J. regia*), which was used as bait, and *Pterocarya rhoifolia* and *P. stenoptera*, which were found at the study site, were excluded from further analysis because their *rbcL* F3R3 sequences were the same. When small mammals, such as *A. speciosus*, are captured by bait traps to estimate plant food resources in their feces, it is necessary to use bait derived from plants or not genetically related plant species that are not growing in the study site.

### Sequence data processing

The *rbcL* F3R3 region was successfully amplified from 100 fecal samples. Sequencing of the *rbcL* F3R3 region yielded a total of 8,895,012 DNA sequences (after quality filtering and removing chimeric sequences) with a mean of 88,950 DNA sequences per fecal sample (S4 Table). In this study, the *rbcL* F3R3 sequence data were normalized to 1,000 DNA sequences, and ASVs were identified using 100,000 DNA sequences. As a result, a total of 201 ASVs were distinguished (S5 Table). All of the nucleotide sequence data were deposited in the DDBJ database (BioProject number PRJDB17374). Based on the results of homology searches using the local *rbcL* F3R3 database, the number of ASVs with more than 98% homology was 185 (92.0%) out of 201 ASVs. For the 16 ASVs (8.0%) that had less than 98% homology using the local *rbcL* F3R3 database, we performed a homology search using the NCBI database. The results showed that 12 out of 16 ASVs were identified as belonging to seven taxa with more than 98% homology using the NCBI (S5 Table). Among these seven plant taxa, *Broussonetia* sp., *Camptotheca acuminate*, and *Croton tiglium*, which are commonly planted for green space maintenance in Japan, were adopted as food plant resources for *A. speciosus.* On the other hand, *Gentiana nipponica, Sieversia pentapetala*, and *Vaccinium* sp., which grow in the alpine meadow zone of Japan, and barley (*H. vulgare* subsp*. Vulgare*), which was the bait used to catch *A. speciosus* in this study, were not adopted as food plant resources for *A. speciosus.* The remaining four ASVs had less than 98% homology with the sequences in our local *rbcL* F3R3 database and those identified based on an NCBI search.

### Identification of food plant resources

The results obtained for the DNA metabarcoding analysis of the food plant sequences are summarized in Table 1. The average number of identified plant taxa per fecal sample was 6.8 plant taxa (median: 6, min: 1, max: 16). The local *rbcL* F3R3 database results identified 192 ASVs with more than 98% homology that could be assigned to 72 plant taxa in 43 families; 43 (59.7%) could be identified to species, 16 (22.2%) to genus, and 13 (18.1%) to family (Table 1). Of the 43 plant families identified using the local *rbcL* F3R3 database, the dominant families throughout all collection periods were Lauraceae (81.0% of 100 fecal samples), followed by Fagaceae (70.0%), Rosaceae (68.0%), and Oleaceae (48.0%). The dominant food plants were *Cinnamomum* sp. (79.0%), followed by Fagaceae-1 (the numbers after the scientific name are identifiers used to distinguish different ASVs within the same taxonomic group; see Table 1 for details) (55.0%), Rosaceae (53.0%), Oleaceae-2 (47.0%), and *Quercus* sp. (39.0%) (Table 1). A total of 50 of 72 plant taxa that were identified as food plant resources were woody plants (Table 1). The predominant food plants detected in more than 20% of the fecal samples in each season are shown in Fig 3. The top four food plant taxa in all seasons were *Cinnamomum* sp., Fagaceae-1, Rosaceae, and Oleaceae-2, and the main food plant resources were the same throughout the year (Fig 3). Comparing the number of food plant taxa in one fecal sample across the four seasons, there were significantly fewer plant taxa in feces sampled in winter than in other seasons (Steel-Dwass test, P<0.05) (Fig 4). The estimated asymptotic Shannon diversity index results for the four seasons indicated a lower diversity of food plants in winter (S_est_ estimated asymptotic Shannon diversity index of 12.6) compared to other seasons (33.2 for spring, 31.4 for summer, and 33.1 for autumn). Rarefaction analysis of the food plant resources consumed in each season at the study site revealed coverage of 86.4% in winter to 93.6% in spring. Furthermore, 96.5% of the food plant taxa during the survey period were covered (Table 2).

1. Database used for homology search (L: rbcL Local database, N: NCBI). For more details, see S5 Table.
2. The sequence of *Melia azedarach* (HE963559) was performed homology search in NCBI and we confirmed misidentified using Blast Tree View. The results showed that *M. azedarach* (HE963559) belonged within the cluster of *Wisteria* sp. Therefore, the sequence of *M. azedarach* (HE963559) was considered misidentified and excluded from this study.
3. The sequence of *Hydrilla verticillata* (KJ747463 - KJ747466) were performed homology search in NCBI and we confirmed misidentified using Blast Tree View. The results showed that *H. verticillata* (KJ747463 - KJ747466) belonged within the cluster of Polygonaceae. Therefore, the sequence of *H. verticillata* (KJ747463 - KJ747466) was considered misidentified and excluded from this study.

**Fig 3.**
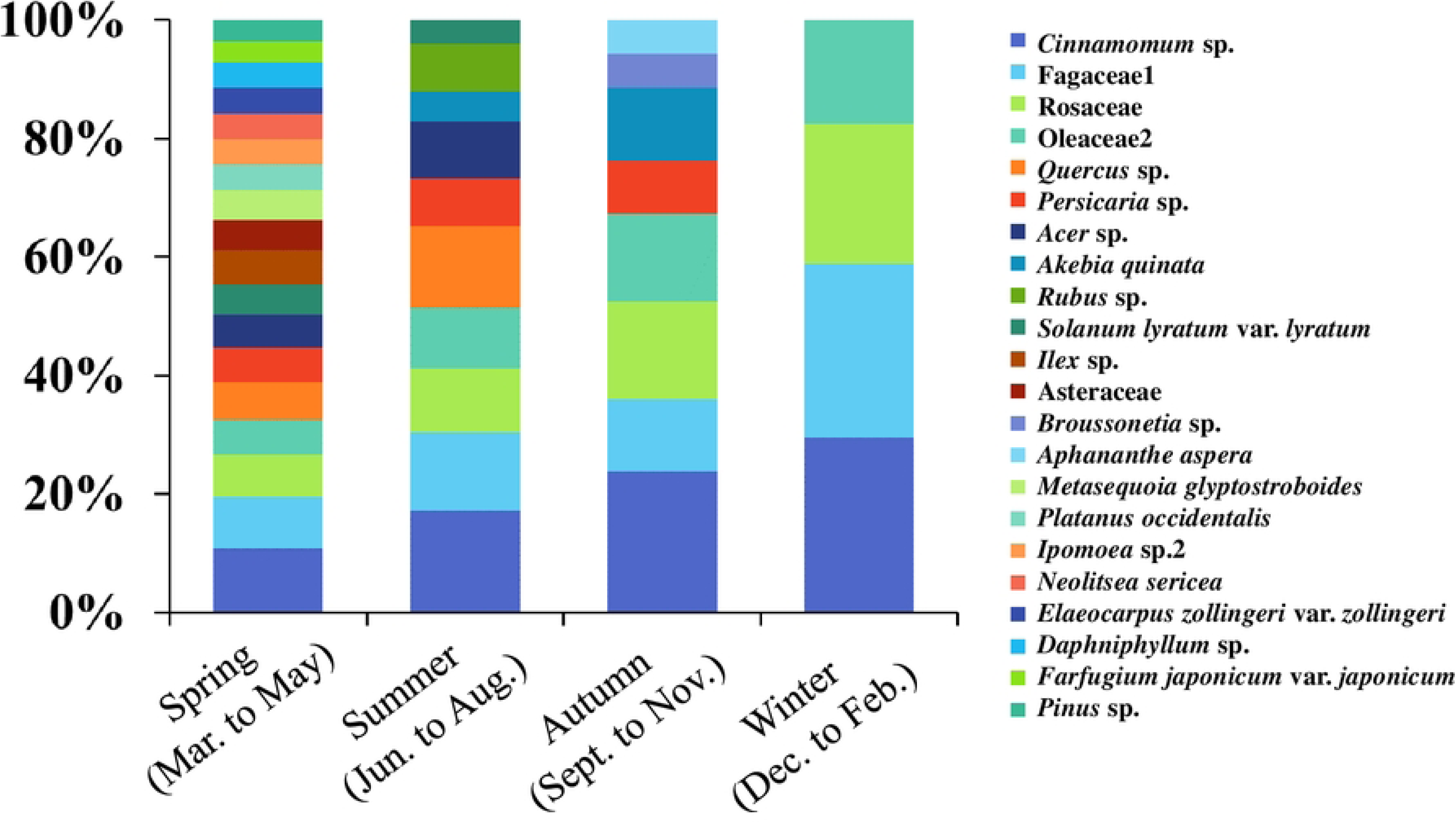
Seasonal changes in dominant food plant taxa. Food plant taxa detected in more than 20% of feces samples in each season.

**Fig 4.**
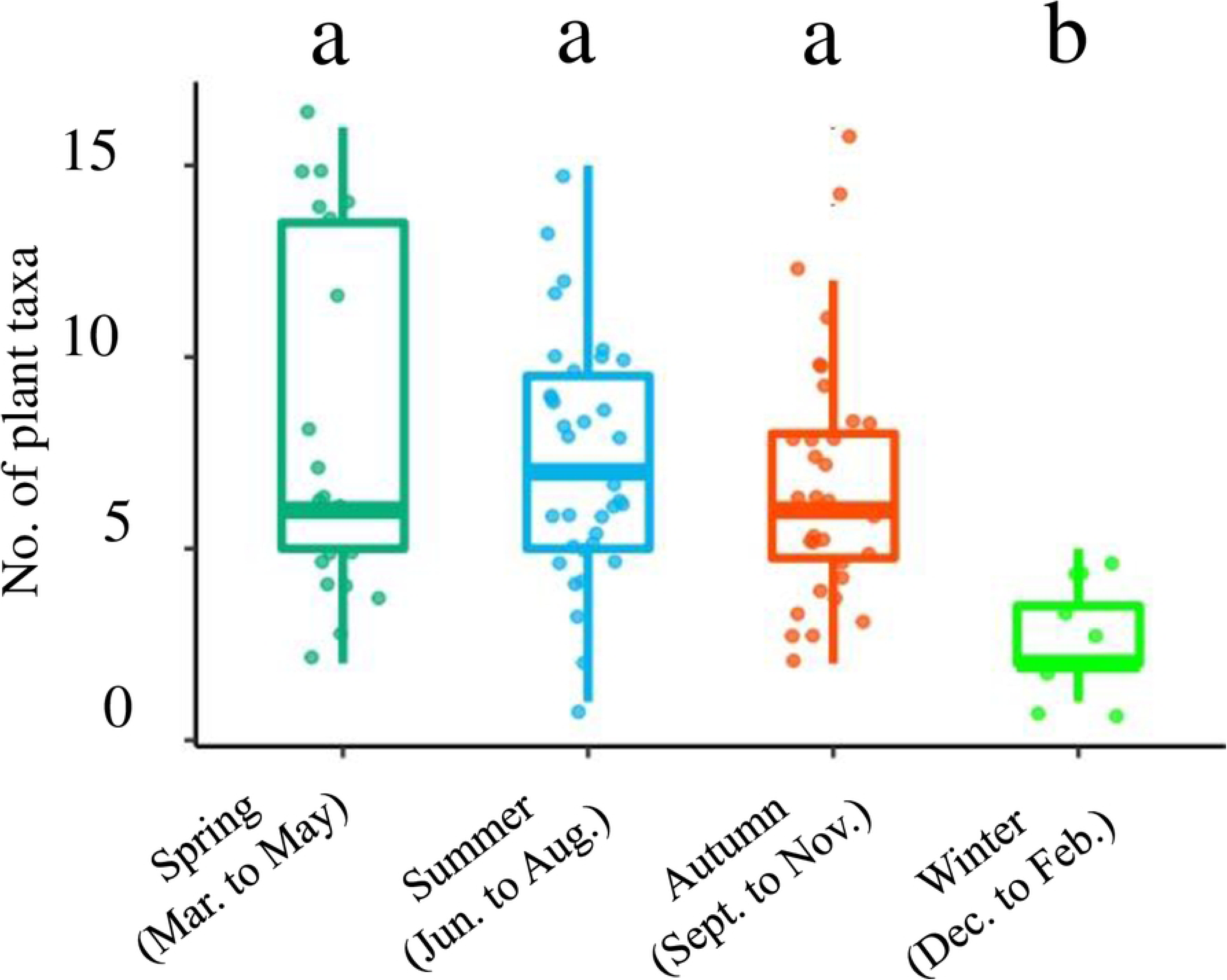
Comparison of the number of plant taxa per fecal sample identified in different seasons.

**Table 1.**
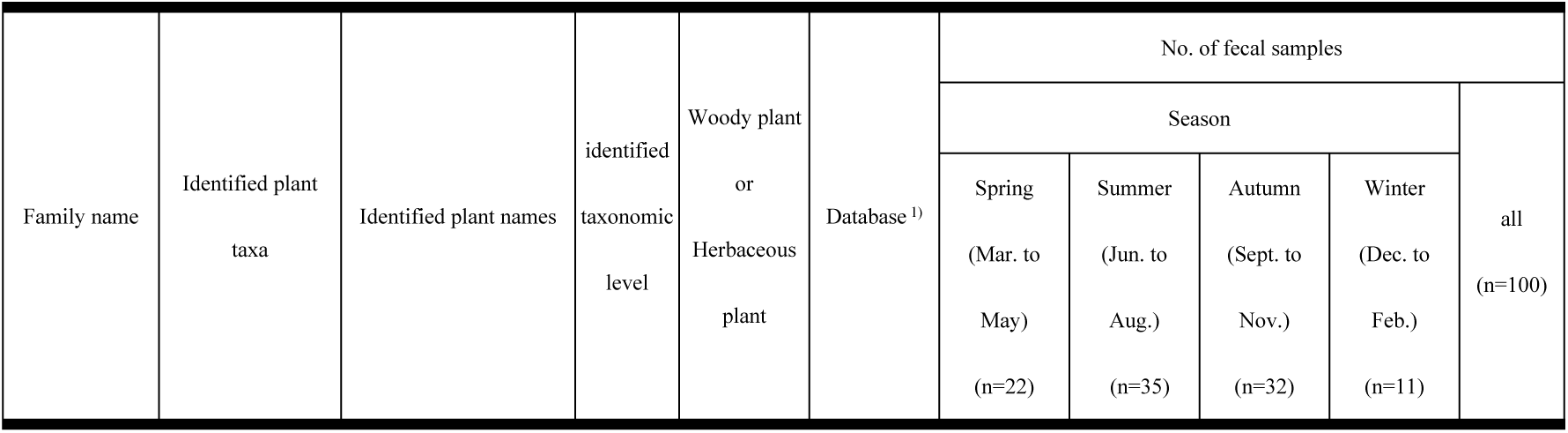
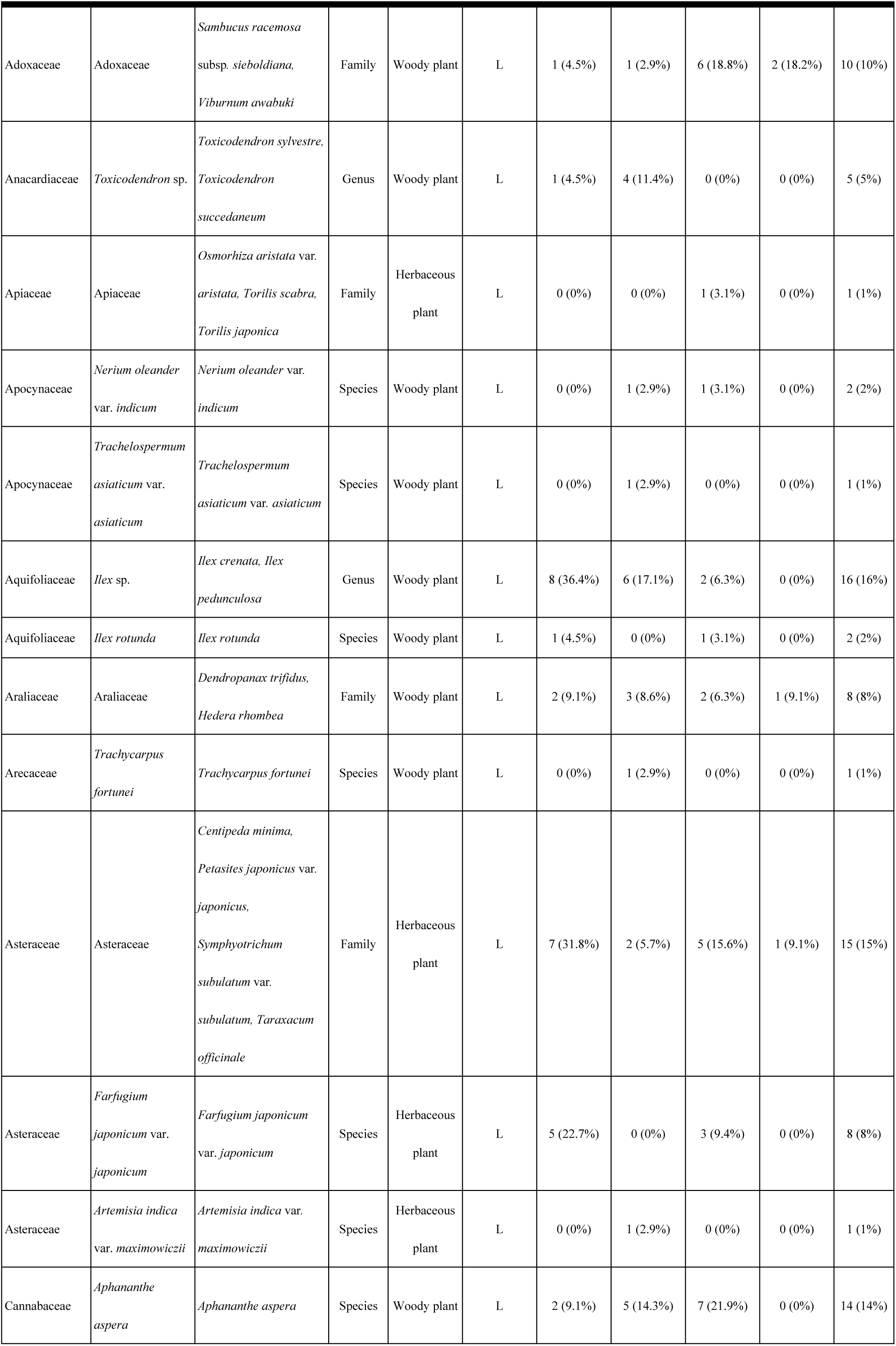
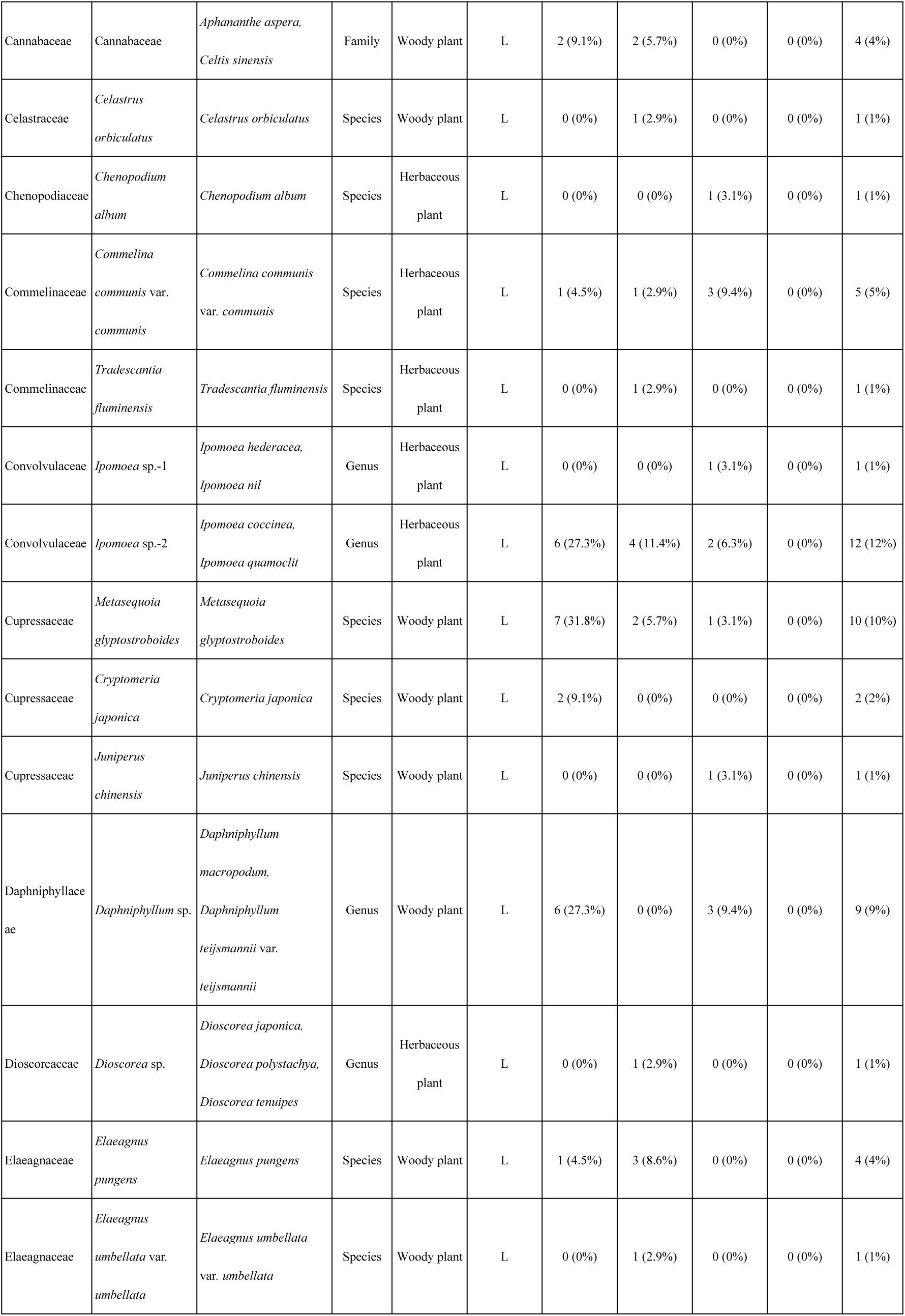
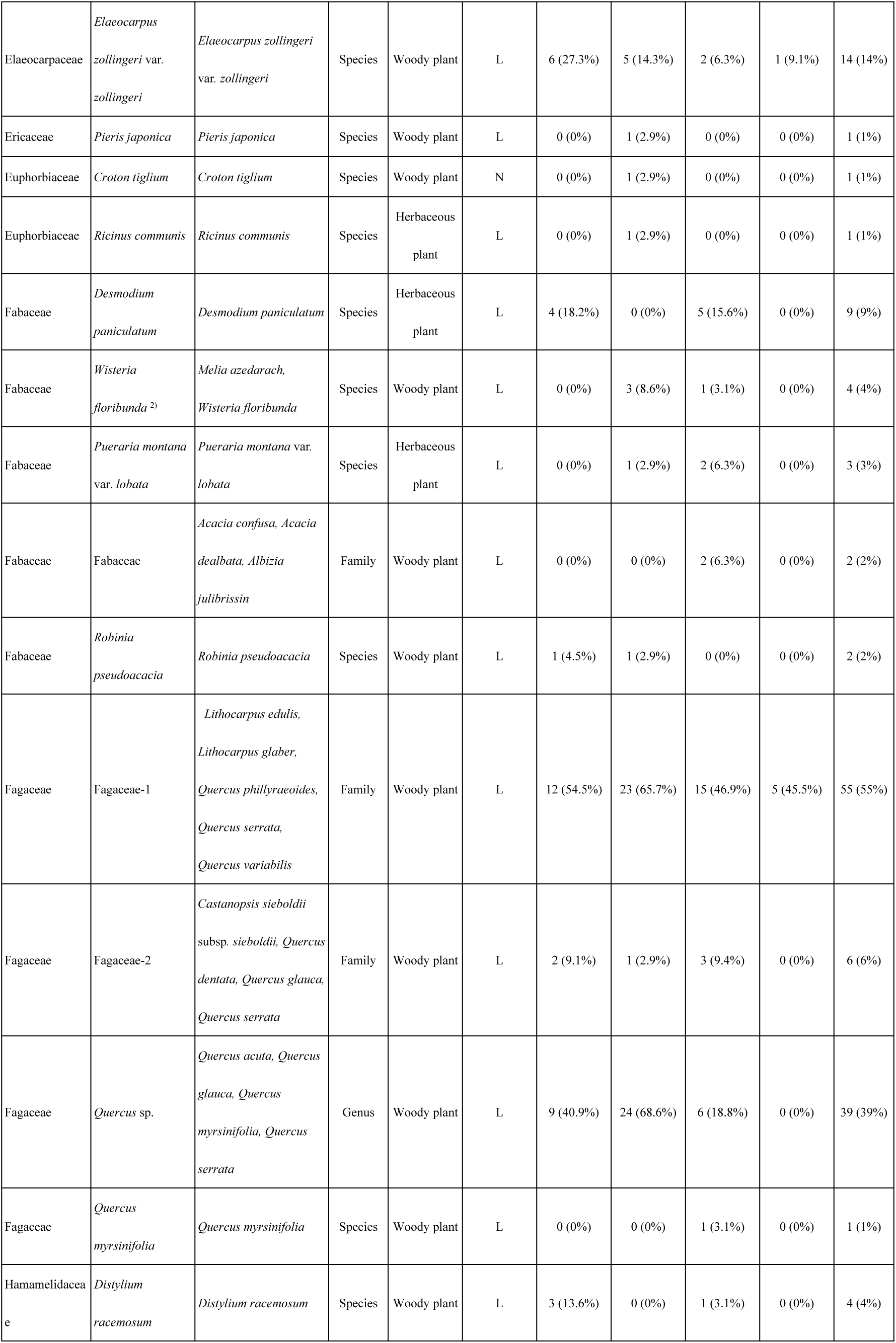
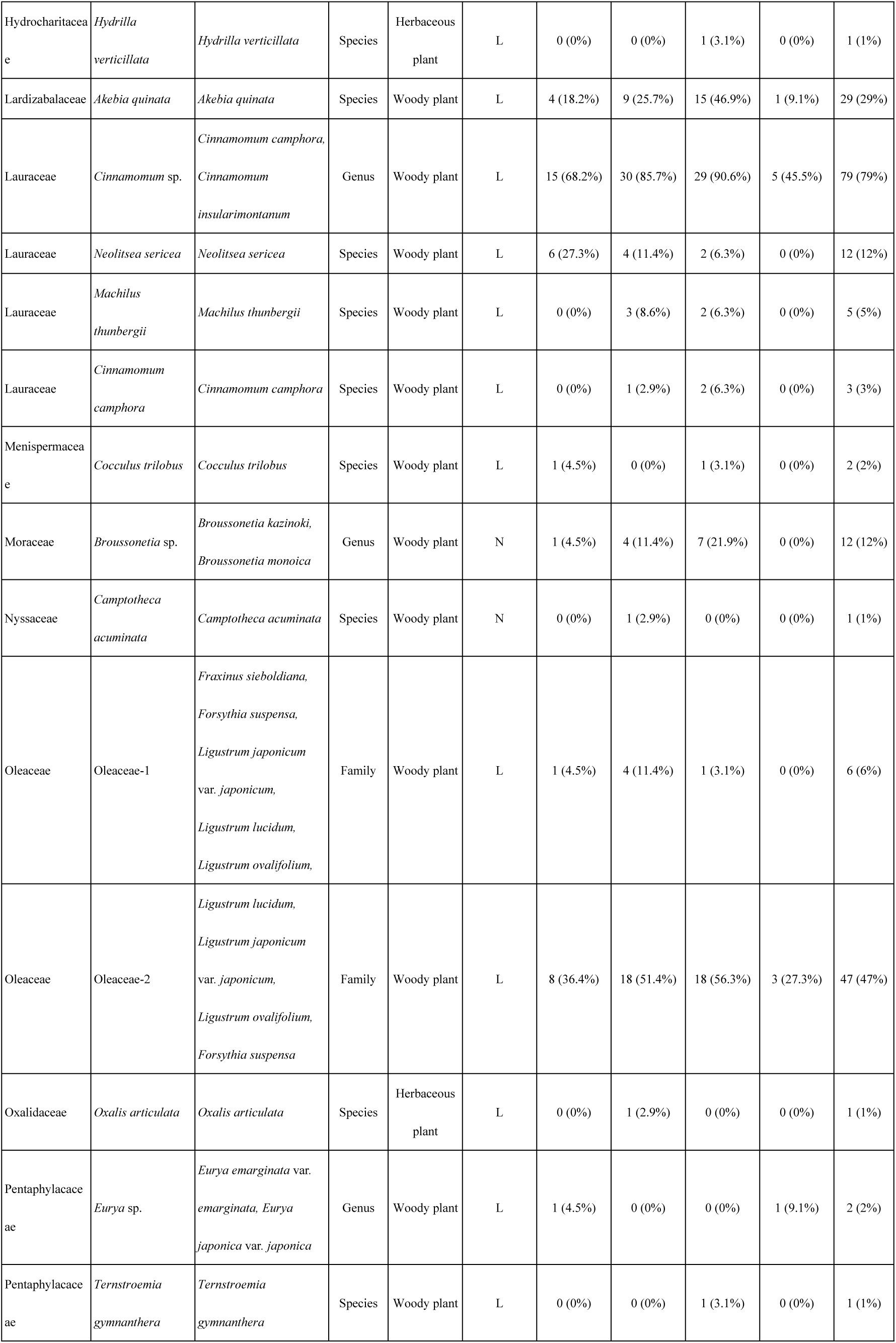
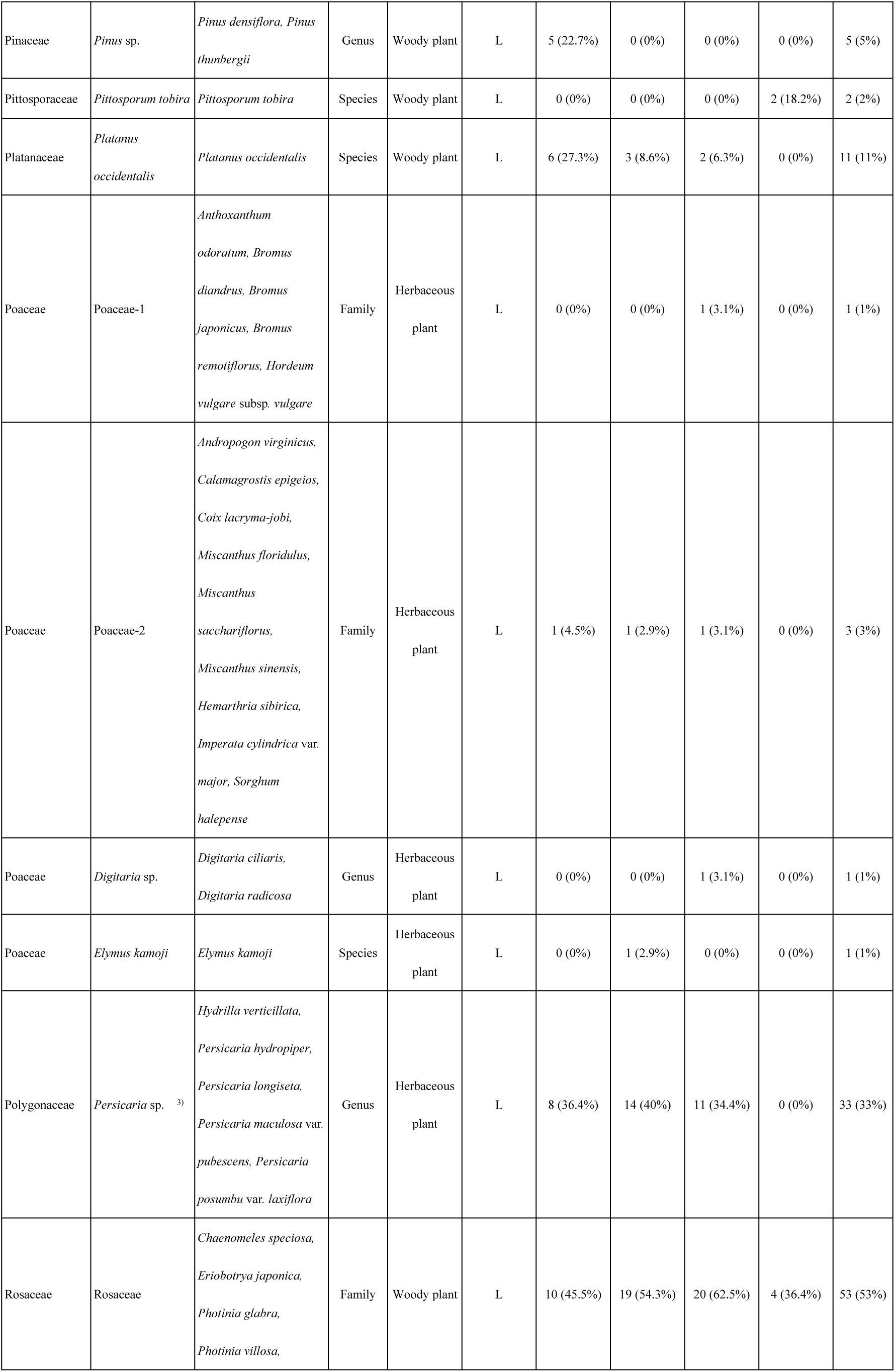
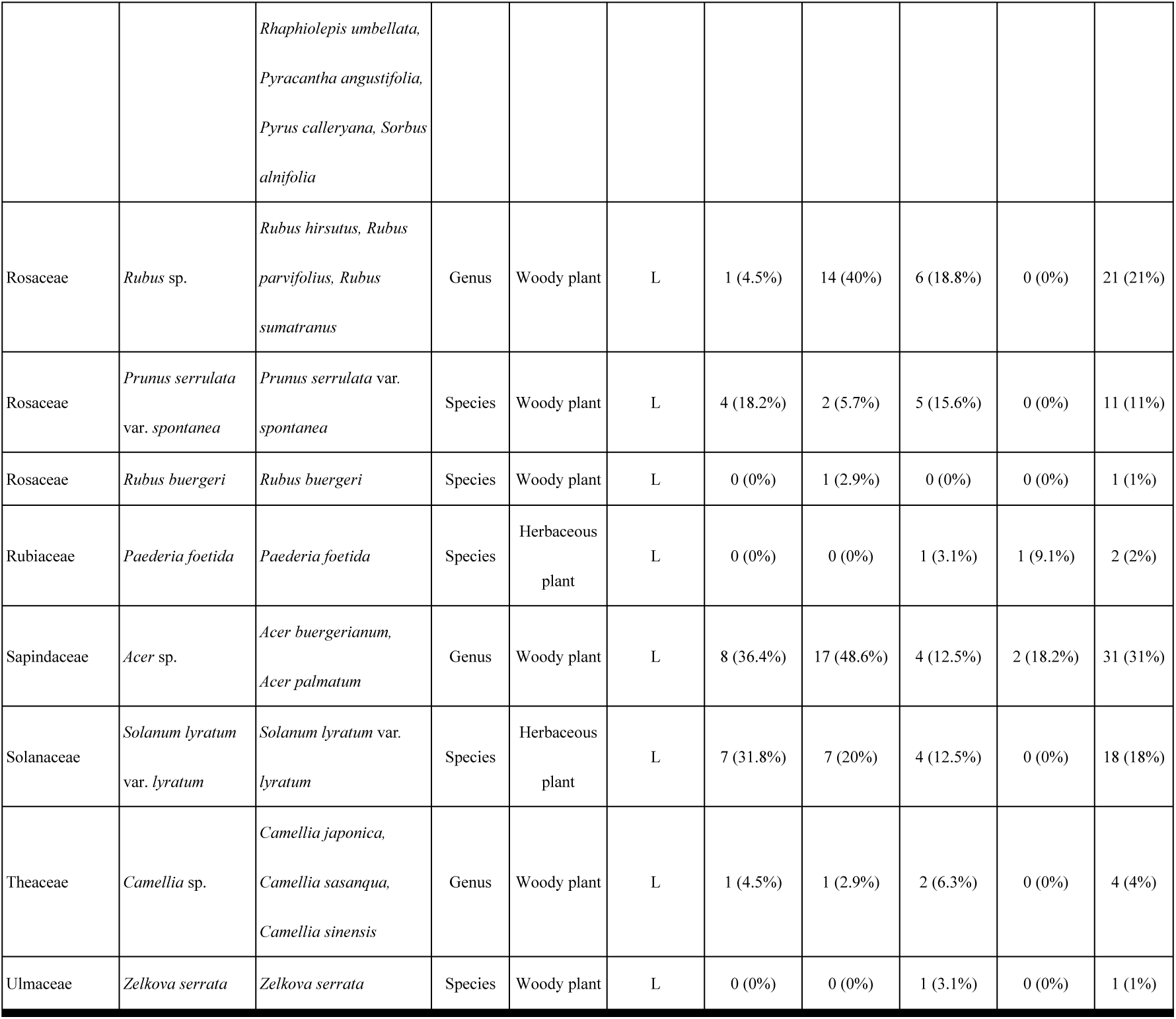
Food plant taxa identified using DNA metabarcoding with rbcL.

**Table 2.** Estimated asymptotic Shannon diversity indexes and coverage of food plant candidates by season.

| Fecal samples collected seasons | No. of fecal samples | No. of identified plant taxa | $S_{obs}$ | $S_{est}$ | Coverage of food plant candidates (%) |
| --- | --- | --- | --- | --- | --- |
| Spring (Mar. to May) | 22 | 41 | 29.0 | 33.2 | 93.6 |
| Summer (Jun. to Aug.) | 35 | 51 | 26.3 | 31.4 | 91.9 |
| Autumn (Sept. to Nov.) | 32 | 51 | 28.6 | 33.1 | 92.1 |
| Winter (Dec. to Feb.) | 11 | 14 | 10.0 | 12.6 | 86.4 |
| Total | 100 | 72 | 32.7 | 35.4 | 96.5 |

$S_{obs}$ observed Shannon diversity index.
$S_{est}$ estimated asymptotic Shannon diversity index.

For each season, the box represents the interquartile range, encompassing values from the 25th to the 75th percentile. The median number of taxa for each season is indicated by the bold line within the box. The ’whiskers’ extend to the most extreme values within 1.5 times the interquartile range from the median. Statistically significant differences between seasons, as determined by the Steel-Dwass test (P<0.05), are denoted by distinct letters.

## Discussion

### Accuracy of food plant resource identification

In this study, the average number of plant taxa identified per fecal sample was 6.8 plant taxa (median: 6, min: 1, max: 16). A total of 72 plant taxa were identified from 100 fecal samples, with 43 (59.7%) assigned to species, 16 (22.2%) to genus, and 13 (18.1%) to family. Rarefaction analysis showed that 96.5% of the food plant taxa during the survey period were covered (Table 2). In previous studies on the food plant resources utilized by *A. speciosus* that were performed by HTS using partial sequences of the chloroplast *trn*L P6 loop intron region without constructing a local database, the average number of plant taxa identified per fecal sample (n=32) in Hokkaido, Japan, was 2.2 (median: 2, min: 1, max: 5) [18]. In that study, a total of 35 plant taxa were identified as food plant resources, with one (2.9%) assigned to one species, 11 (31.4%) to genus, and 23 (65.7%) to family. On islands in the Seto Inland Sea in western Japan, the average number of plant taxa identified per *A. speciosus* fecal sample (n=76) was 4.3 (median: 4, min: 1, max: 14), with a total of 70 plant taxa identified, which could be assigned to 11 species (15.7%), 22 genera (31.4%), 36 family (51.4%), and one taxon that could not be identified to family level (less than 98% homology) [19]. Although the survey site, survey period, and DNA region sequenced by HTS differed between the present study and the aforementioned reports [18, 19], both the accuracy of plant species identification and the number of plant taxa detected per fecal sample in this study were higher than those of the previous studies. In the present study, 60 mg of the feces excreted in the trap by a single individual were analyzed, whereas in the previous studies [18, 19], only three fecal pellets (weight not reported, but likely less than 60 mg) excreted in the traps by one individual were analyzed; this may have led to an underestimation of the food resources foraged by one individual. In addition, the possible reason for the relatively higher accuracy observed in this study may be because the previous studies did not construct a local *trn*L P6 loop intron region database for DNA metabarcoding the plants found at their study site.

### Dominant food plant resources

In the Japanese Archipelago, a major component of the diet of *A. speciosu*s is acorn-producing *Quercus* species [17–19, 52]. However, in this study, materials from the family Lauraceae (81.0% of all 100 fecal samples) were dominant in fecal samples, with *Cinnamomum* sp. (79.0%) (presumed to be either *C. camphora* or *C. insularimontanum* from our vegetation survey) also identified as a dominant food plant resource for the *A. speciosus* population at this study site throughout the year. Although *Cinnamomum* sp. have been reported to be utilized as a food resource by *A. speciosus*, to the best of our knowledge, there are no previous records of this genus being a dominant food plant source for *A. speciosus* [17–19, 26, 52–54]. *Apodemus speciosus* tends to specialize in consuming acorns from *Quercus* species [16, 18, 19]. However, in this study area, *Cinnamomum* species were more dominant in the feces samples than *Quercus* species, suggesting that *Cinnamomum* species were the primary food resource of *A. speciosus* at the study site. The subdominant families in the feces samples included Fagaceae (70.0%) (mostly *Quercus* sp.), Rosaceae (68.0%), and Oleaceae (48.0%) (presumed to be *Ligustrum* sp. from our vegetation survey). The results are consistent with previous reports which showed that *A. speciosus* utilizes a variety of food plants, and seeds in particular [17–19, 26, 53, 55]. Since there were few acorn-bearing *Quercus* trees at this study site, it is possible that the seeds of *Cinnamomum* sp., Rosaceae, Oleaceae-2 are used in winter (Table 1). The herbaceous plants that were utilized by *A. speciosus* at the study site were dominated by *Persicaria* sp. (33.0%), followed by *Solanum lyratum* var. *lyratum* (18.0%) and members of the Asteraceae (15.0%) (Table 1). The *A. speciosus* population at the study area fed more on woody plants than on herbaceous plants, which is consistent with the conclusions of previous studies [18, 19]. The planting and maintenance of woody plants is therefore important for maintaining the population of *A. speciosus* in the study area (Table 1).

As shown in Fig 3, the main food plant resources utilized by the *A. speciosus* population in our study area did not differ among seasons. In winter, food plants are scarce; however, the dominant food plant resources are the same as in other seasons. On the other hand, the number of plant taxa identified per single sample was significantly lower in winter than in spring, summer, and fall (Steel-Dwass test, P<0.05) (Fig 4). The estimated Shannon diversity value was also lowest in winter (12.6), less than half that recorded in the other seasons. This decrease in the diversity of food plant resources in winter is likely due to the fact that fewer food plant resources are available in winter. In order to develop artificial green spaces in the study area in a way that considers the habitat suitability of *A. speciosus*, it is considered necessary to plant and maintain not only representatives of *Cinnamomum* sp., Fagaceae (mostly *Quercus* sp.), Rosaceae, and Oleaceae, which provide food resources for *A. speciosus* throughout the year, but also *Pittosporum tobira*, which is foraged frequently in winter. In addition, planting of acorn-producing *Quercus* species, should also be considered.

## Conclusions

In conclusion, the DNA metabarcoding analysis described in this study provides a better understanding of the feeding habits of *A. speciosus* inhabiting artificial green spaces. In this study, in order to construct the local *rbcL* F3R3 database, the *rbcL* F3R3 sequences of plant species that could not be collected in the study area were obtained from the NCBI database. As a result, the findings presented here are considered to accurately reflect the food plant resources utilized by the Large Japanese field mouse *A. speciosus* living in artificial green spaces. Therefore, these food plant resource data can contribute to the management of artificial green spaces in Japan with a focus on biodiversity conservation.

## Acknowledgements

We thank the staff of the Aichi Complex of Idemitsu Kosan Co., Ltd., Chita Thermal Power Station of JERA Co., Inc. (formerly Chita Thermal Power Station of Chubu Electric Power Co., Ltd.), and ENEOS Corporation for access to the study area. We thank Mr. Kitou Nobuyuki and Ms. Shibata Akane of Chubu University for their assistance with capturing mice.

## Supporting information

**S1 Table. List of initial PCR primers**

List of primers used for the initial PCR. Letters in italics indicate the MiSeq sequencing primers. Bold Ns indicate random bases used to improve the quality of MiSeq sequencing. Single underlined letters indicate DNA barcoding primer sequences.

**S2 Table. List of second PCR primers**

List of primers used for the second PCR. Letters in italics indicate the MiSeq sequencing primers. Bold Xs indicate index sequences used to identify each sample. Double-underlined characters indicate P5/P7 adapter sequences for MiSeq sequencing.

**S3 Table. Local *rbcL* F3R3 database**

**S4 Table. Sequencing statistics of *rbcL* regions**

Input reads: number of reads in raw fastq files.

Filtered reads: number of reads after preliminary quality filtering. DenoisedF/R reads: number of reads after quality filtering.

Merged reads: number of merged forward-reverse reads.

Nonchim reads: number of merged reads after removal of chimeric sequences.

**S5 Table. Results of homology searches for fecal samples using the local *rbcL* database and the NCBI database**

